# CPIExtract: A software package to collect and harmonize small molecule and protein interactions

**DOI:** 10.1101/2024.07.03.601957

**Authors:** Michael Sebek, Andrea Piras, Lucía Prieto-Santamaría, Shi Chenghao, Gordana Ispirova, Giulia Menichetti

## Abstract

**Summary:** The binding interactions between small molecules and proteins are the basis of cellular functions. Yet, experimental data available regarding compound-protein interaction are not harmonized into a single entity but rather scattered across multiple institutions, each maintaining databases with different formats. Extracting information from these multiple sources remains challenging due to data heterogeneity. Here, we present CPIExtract (Compound-Protein Interaction Extract), a tool to interactively extract experimental binding interaction data from multiple databases, perform filtering, and harmonize the resulting information, thus providing a gain of compound-protein interaction data. When compared to a single source, DrugBank, we show that it can collect more than ten times the number of annotations. The end-user can apply custom filtering to the aggregated output data and save it in any generic tabular file suitable for further downstream tasks such as network medicine analyses for drug repurposing and cross-validation of deep learning models.

**Availability:** CPIExtract is an open-source Python package under an MIT license. CPIExtract can be installed from https://pypi.org/project/cpiextract and downloaded from https://github.com/menicgiulia/CPIExtract. The package can run on any standard desktop computer or computing cluster.

## Introduction

Compound-protein interactions (CPI) dictate the basis of many cellular dynamics, such as signaling, enzymatic reactions, and gene regulation. By understanding these interactions, we can develop drugs to target specific disease phenotypes and study how dietary molecules affect human and microbial health. Often CPI information is generated with high-throughput screening, testing small molecules (≤ 1000 Daltons) against large panels of tissues, cells, or proteins to determine their bioactivity. The generated data are gathered and curated into multiple chemical data sources such as BindingDB (Gilson et al., 2016), ChEMBL (Bento et al., 2014), and PubChem (Kim et al., 2023). Each database is managed by different organizations, each implementing its own format, resulting in significant heterogeneity in the types of information stored. For example, databases with a focus on pharmaceuticals such as DrugBank (Knox et al., 2024) and DrugCentral (Avram et al., 2023) mainly focus on reporting therapeutic targets, whereas databases like STITCH (Szklarczyk et al., 2016) and the Comparative Toxicogenomics Database (Davis et al., 2021), despite compiling extensive biological interaction networks, lack in-depth information on the bioactivities of these interactions. The presence of crowd-sourced databases like Drug Target Commons (Tanoli et al., 2018) and Open Targets Platform (Ochoa et al., 2023) further exacerbates this diversity as each user follows individual reporting criteria. Consequently, extracting and integrating CPI data from these sources poses a significant challenge to researchers. The importance of interaction data has driven efforts to unify chemical structure notation (Pascazio et al., 2023), protein names (Seal et al., 2023), and protein-protein interaction data (Dimitrakopoulos et al., 2021). Yet, there is currently no solution for consolidating CPI data despite being essential to many applications, including the training of AI pipelines for drug-target discovery (Chatterjee et al., 2023), developing chemical language models for de-novo drug design (Grisoni, 2023), and serving as seed interactions in network medicine analyses (Nasirian and Menichetti, 2023). The sparsity of CPI data leads researchers to develop predictive tools for annotating understudied compounds and proteins (Tsubaki et al., 2019). However, the performance and generalizability of these models are hindered by the extensive effort needed to integrate different data sources. To address these challenges, we present CPIExtract, a Python package enabling users to seamlessly extract experimental interaction data from nine compound-protein interaction databases. By automating the filtration, integration, and standardization processes, CPIExtract generates a unified output that can be stored in any standard tabular file format.

## CPIExtract Pipeline Workflow

The package allows the user to execute two pipelines, Compound-Proteins-Extraction (Comp2Prot) and the Protein-Compounds-Extraction (Prot2Comp). The Comp2Prot pipeline retrieves proteins interacting with the small molecule passed as input, while Prot2Comp returns compounds interacting with the input protein. Both pipelines comprise three phases: input data extraction, data filtering, and harmonization (Fig. 1). The package is designed to support multiple user scenarios based on different storage and software availability (SI User scenarios).

**Figure 1.**
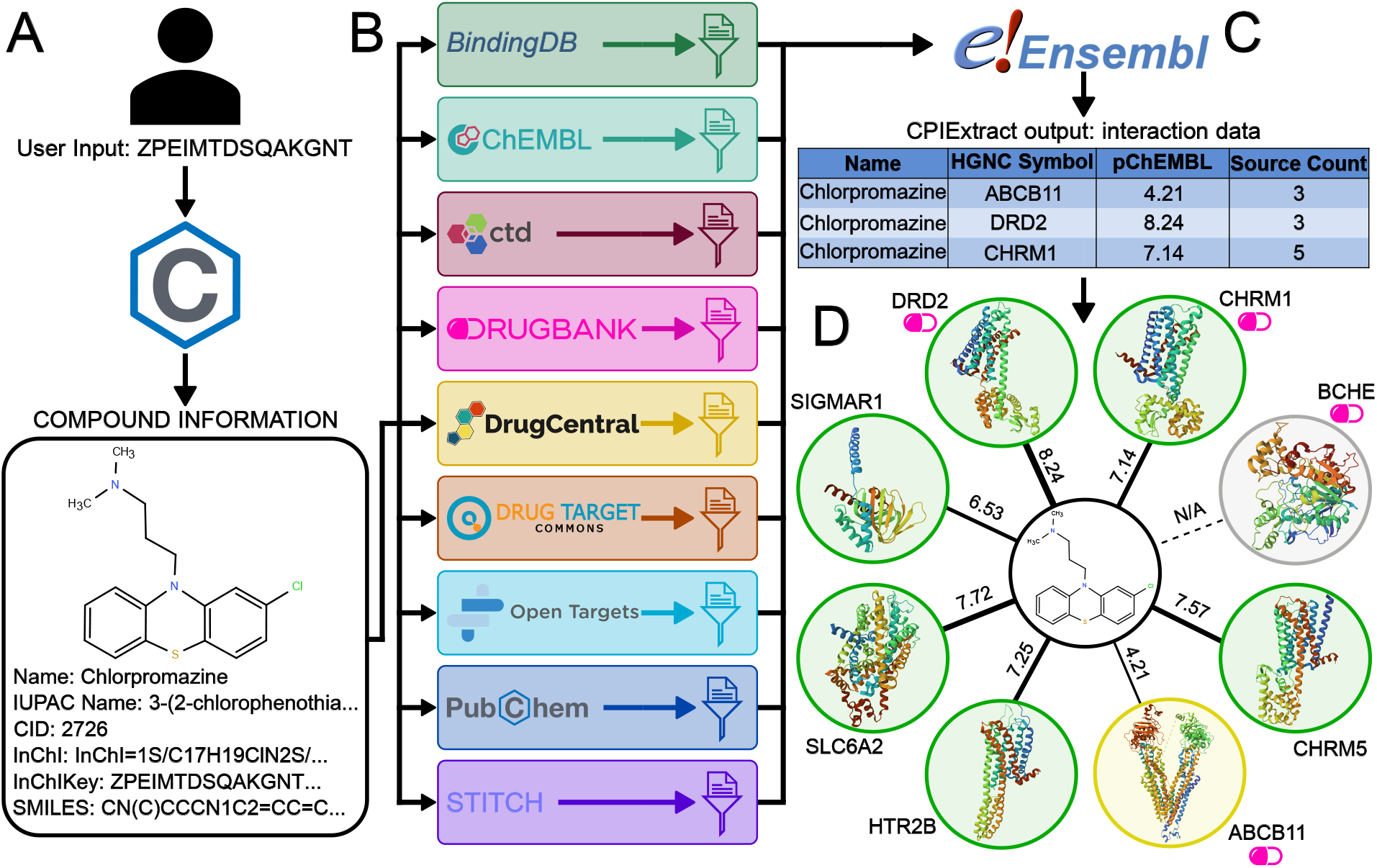
CPIExtract pipeline. Schematic of the Comp2Prot pipeline to extract CPI data for a compound. **A)** User provides a compound identifier which is used to query PubChem. **B)** CPIExtract collects and filters interaction data for each database, and **C)** standardizes the protein identifiers via Ensembl, resulting in a tabular output. **D)** Interaction network of Chlorpromazine CPI data (green: strong binding, yellow: weak binding, gray: no data). The DB logo marks annotations in DB.

### Input data extraction

CPIExtract extracts raw data from the following nine databases: BindingDB (BDB), ChEMBL, Comparative Toxi-cogenomics Database (CTD), DrugBank (DB), DrugCentral (DC), Drug Target Commons (DTC), Open Targets Platform (OTP), PubChem, and STITCH. Comp2Prot accepts InChI, InChIKey, SMILES, and PubChem ID (CID) as input, while Prot2Comp accepts Entrezgene ID, HGNC ID, Ensembl Peptide ID, Ensembl Gene ID, UniProt ID, ChEMBL ID, and HGNC symbol as input. To unify databases and remove duplicates, compound identifiers are resolved via the PubChem API (PubChemPy Python package) while protein annotations are harmonized using either the Ensembl BioMart API (biomart Python package) or MyGene (mygene Python package).

### Data filtering

CPIExtract focuses exclusively on *Homo sapiens* proteins and protein interactions. Thus, the package removes non-human data and eliminates non-protein interactions, such as RNA interactions, mutated protein interactions, and protein complex interactions. Afterwards, the pipeline curates each database depending on the available data: 1) Activity Description: removes interactions with statements indicating inconclusive or undetermined activity (BDB, ChEMBL, CTD, DTC, PubChem); 2) Bioactivity Measurements: removes interactions not reporting binding activity (BDB, ChEMBL, DC, DTC, PubChem); 3) Reporting Source: filters to ensure interactions have at least one literature source (ChEMBL, DTC, OTP) or at least one reported experiment, removing all text mining and predicted evidence (STITCH); 4) Statements of confirmed binding (CTD), for example CTD has labels like ‘affects^binding’; 5) Experimental Conditions: filters interactions by temperature and pH (BDB), and use descriptions of condition validity (ChEMBL), and 6) documented mechanisms of action (OTP). Experimental conditions reflect physiological ranges across human tissues and subcellular compartments: temperature (< 40°C, the clinical threshold for hyperthermia (Laupland, 2009)) and pH (5 < pH < 9, encompassing values from acidic endolysosomal compartments to alkaline secretions (Casey et al., 2010)). Since these data are typically only reported when deviating from expected conditions, null values are assumed to represent standard conditions. The package supports optional filters (SI Optional Filtering Parameters).

### Data harmonization

CPIExtract harmonizes the collected bioactivity data by calculating the pChEMBL value, which is defined as -*log*_10_(molar IC50, XC50, EC50, AC50, *K*_*i*_, *K*_*d*_, or potency) and provides a numerical value for the binding strength (Bento et al., 2014). BDB, ChEMBL, DC, and PubChem report bioactivity using these standardized metrics, while the user-submitted data in DTC are converted to standardized metrics whenever possible. CPIExtract calculates the average pChEMBL value for unique bioactivities, as databases frequently reference the same sources. We acknowledge that IC50, *K*_*i*_, *K*_*d*_, and EC50 values are not directly interchangeable. Yet, drug discovery pipelines largely employ activity thresholds (non-binding pChEMBL ≤ 3, weakly binding 3 *<*pChEMBL*<* 6, and strongly binding pChEMBL ≥ 6) where cross-assay variability for a given compound–protein pair is unlikely to produce systematic misclassifications (Chatterjee et al., 2023, Kramer et al., 2012). The results are provided to the user in a tabular format, which includes the compound and protein identifiers, interaction strength via pChEMBL, and the databases reporting each interaction.

## Applications

To prove the effectiveness of CPIExtract, we compare it to DrugBank (DB), one of the most important drug-target databases, widely used as training data for machine learning models (Tsubaki et al., 2019) and as a data source for network medicine applications (Sadegh et al., 2021). DB currently contains more than 16,000 drugs and 5,700 proteins with more than 35,000 compound-protein interactions for *Homo sapiens*. When focusing on small molecule drugs, DB has 11,340 reported compounds; however, only 5,478 of these compounds have reported interactions (Fig. 2A). Comp2Prot finds interaction data for 8,221 compounds, an additional 2,743 compounds more than DB (50.1% more coverage). In addition, it collects 219,715 interactions compared to the 21,128 interactions in DB, corresponding to a 10-fold increase. Lastly, the interactions found by CPIExtract link the compounds to a total of 9,581 proteins, improving the coverage from the 2,745 proteins in DB. The significant increase in CPI data with CPIExtract reveals a fundamental bias in research databases, which often focus on compounds and proteins with known health implications. CPIExtract addresses this bias by integrating diverse datasets, aiding in the mapping of crucial off-target effects.

**Figure 2.**
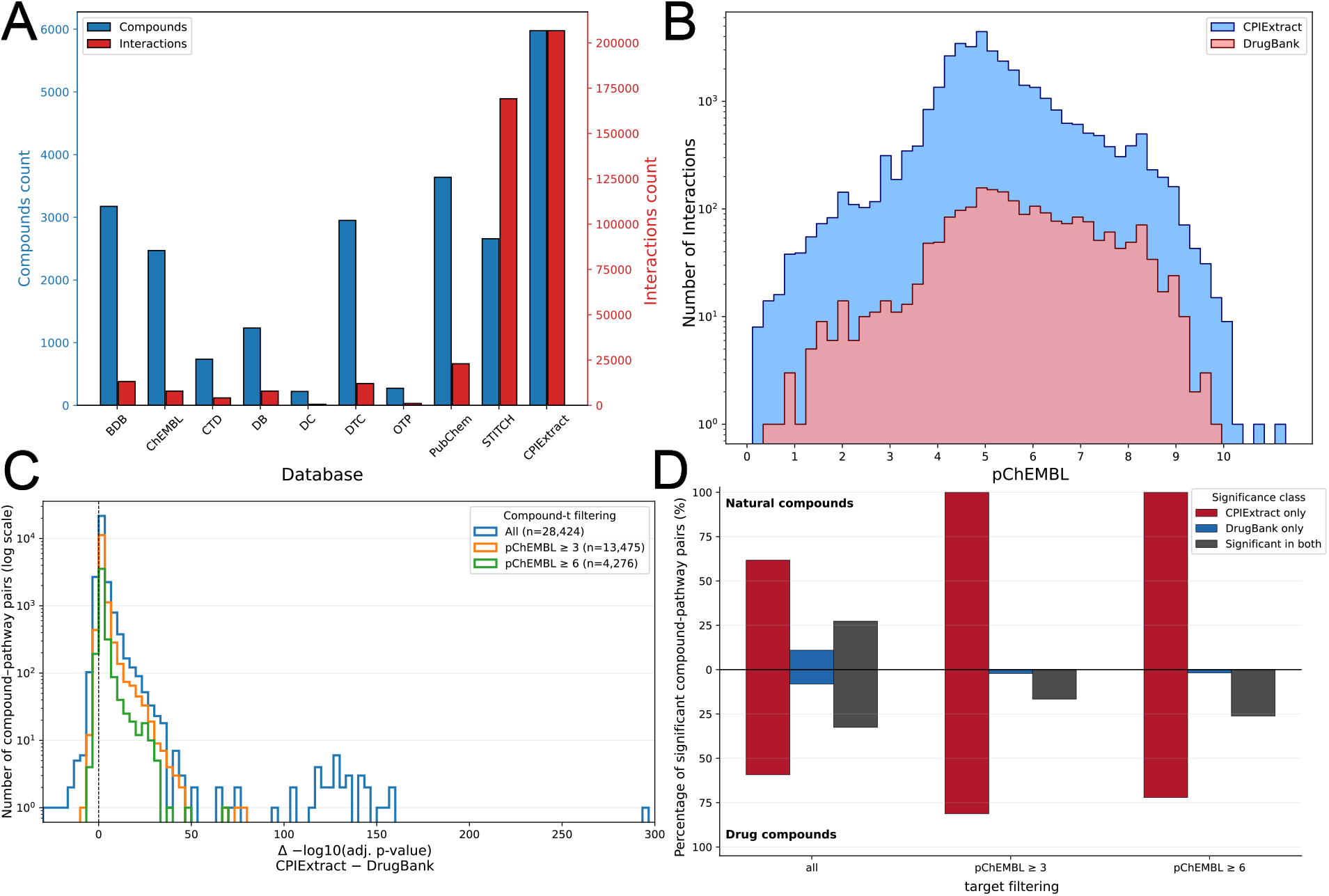
CPIExtract-DrugBank Comparison. **A)** CPI data found in each database for the small molecule drugs within DB (blue: number of compounds, red: number of interactions). **B)** The pChEMBL distribution for interactions in CPIExtract and DB. **C)** Distribution of △(log_10_(FDR)) between CPIExtract and DB for compound-pathway pairs (blue: all interactions, orange: pChEMBL ≥ 3, green: pChEMBL ≥ 6). Dashed vertical line denotes no difference. Positive values indicate pathways that are more significantly enriched using CPIExtract than DrugBank, whereas negative values indicate stronger enrichment with DrugBank. **D)** Percentage of significant compound-pathway pairs in CPIExtract only (red), DrugBank only (blue), or both resources (gray). Natural compounds are defined as those carrying natural product ATC codes in DB.

Next, we consider the strength of the interactions within the DB and CPIExtract data. We use pChEMBL as the evaluation metric, which we divide into three categories: non-binding (pChEMBL ≤ 3), weakly binding (3 *<*pChEMBL*<* 6), and strongly binding (pChEMBL ≥ 6). While DB does not report bioactivity, CPIExtract can find bioactivity for 29.1% of the interactions, resulting in 0.5 % non-binding, 10.8 % weakly binding, and 17.8 % strongly binding interactions. With CPIExtract, we observed a remarkable 21-fold increase in weakly binding interactions and a 5.1-fold increase in strong interactions (Fig. 2B). Overall, CPIExtract retrieved 5 to 10 times more interactions for each pChEMBL bin value compared to DB, facilitating contrastive learning strategies (Singh et al., 2023) and providing valuable examples of experimentally validated negatives to improve the training of binary binding predictors (Chatterjee et al., 2023). Users can filter the retrieved interactions based on the desired pChEMBL value or the number of sources validating the interaction. We conducted similar analyses for DB proteins (see SI Prot2Comp on DB) and food compounds (see SI Comp2Prot on Naturally Occurring Compounds).

To further assess whether this increase in CPI data from CPIExtract reflects biological meaning, we performed pathway enrichment analysis across the three pChEMBL thresholds. For each compound, we tested for significant enrichment of KEGG pathways in the reported protein interactions from each source, retaining all pathway-compound associations meeting statistical significance (FDR-adjusted p*<*0.05) in either CPIExtract or DB. We then compared both resources by calculating the difference in pathway significance as Δ(−log_10_(FDR)), where positive values indicate stronger enrichment with CPIExtract (Fig. 2C). The distributions are consistently shifted towards positive Δ(− log_10_(FDR)) values across all filtering thresholds (medians larger than 2 among nonzero differences; Wilcoxon signed-rank test, p < 0.001 for all thresholds), indicating that KEGG enrichment obtained with CPIExtract is generally more significant than those obtained with DrugBank. This indicates that the expanded CPI data translates into improved pathway-level resolution. When we stratify the compounds by their ATC codes into natural and drug compounds (SI KEGG pathway enrichment), we find that the gain in pathway enrichment is more prominent for natural compounds (Fig. 2D). CPIExtract uniquely recovers 100% of the significant compound–pathway pairs for natural compounds at both weak- and strong-binding thresholds. It also identifies 81% and 72% of the significant pairs for drug compounds at the weak- and strong-binding thresholds, respectively, whereas DB uniquely recovers only 2% of these pairs. When all compound–pathway pairs are considered, CPIExtract continues to outperform DB, capturing 89% of the significant pairs for natural compounds compared with 38% recovered by DB and 92% of significant pairs for drug compounds in comparison to 41% by DB. These results indicate that the additional interactions recovered by CPIExtract are neither redundant nor dominated by noise; rather, they capture biologically meaningful signals, particularly for naturally occurring compounds whose bioactivity remains substantially underrepresented in curated databases.

## Discussion

CPIExtract offers an extensive and integrated resource for protein-ligand binding interactions, addressing challenges across various domains by merging heterogeneous data and mitigating bias and annotation imbalances from single sources. Specifically, CPIExtract facilitates the stratification of binding information by chemical classes, enabling the creation of balanced datasets that can enhance model training performance. In the field of network medicine, CPIExtract’s comprehensive interaction lists enhance the prediction of health implications (Aldana et al. [2025]). In food science, the tool addresses the issue of limited bioactivity knowledge across databases, providing a more robust framework for understanding compound interactions. Overall, CPIExtract empowers researchers to thoroughly characterize their compounds or proteins of interest, systematically exploring mechanisms of action with robust experimental evidence.

## Author Contributions

M.S., A.P., and L.P-S. developed the methodology. M.S., A.P., S.C., and G.I. built the software. M.S., A.P., L.P-S., and G.M. contributed to the writing and editing. G.M. conceptualized, administered, and funded the project.

