## Supplementary material for "CPIExtract: A software package to collect and harmonize small molecule and protein interactions": SI

### CPIExtract: Supplementary Information

#### Table of Contents


**How to run the tool** **2**

**User scenarios** **3**

Local execution . . . . 4

Server Execution . . . . 4

**APIs limitations and runtime performance** **5**

**Database Information at Publication** **6**

**Database selection** **10**

**Optional Filtering Parameters** **11**

**Using CPIExtract in Computational Approaches** **12**

**CPIExtract on Food Compounds** **14**

**KEGG pathway Enrichment** **16**

**Comp2Prot on DB** **24**

DrugBank Candidates Selection for Evaluation . . . . 25

**Prot2Comp on DB** **27**

**Comp2Prot execution on Naturally Occurring Compounds** **31**

### How to run the tool

CPIExtract is available on the Python Package Index (PyPI) and can be installed via:

```
pip install cpiextract
```

The tool provides two Python classes: `Comp2Prot` and `Prot2Comp`. These pipelines can be executed either locally or using a MySQL server connection, specified via the `execution_mode` argument (`'local'` or `'server'`). In local mode, users pass a dictionary of `pandas` DataFrames via the `dbs` argument. In server mode, users provide connection configuration (IP address, user, password, and database name). Once instantiated, pipelines are run via `comp_interactions()` or `comp_interactions_select()` for Comp2Prot and for Prot2Comp use `prot_interactions()` or `prot_interactions_select()`. The `interactions` functions will run all databases within CPIExtract while the `interactions_select` functions will only run a given set of databases given by the user. To query selected databases only, `interactions_select()` functions accept a `selected_dbs` argument as an underscore-separated string (e.g., `'pc_chembl_db'`; the full string `'pc_chembl_bdb_stitch_ctd_dtc_otp_dc_db'` equates to querying all databases).

The following minimal example demonstrates a complete Comp2Prot query using server mode:

**Option: Server mode**

```
from cpiextract import Comp2Prot

C2P = Comp2Prot(execution_mode='server',
                server_info={"host": "XXX.XX.XX.XXX",
                             "user": "user",
                             "password": "password",
                             "database": "cpie"})

# Chlorpromazine (InChIKey)
```

```
[interactions_agg, db_states, interactions_total]= C2P.comp_interactions(
    input_id='ZPEIMTDSQAKGNT-UHFFFAOYSA-N',
    pChEMBL_thres=3.0, strong_positive_thres=6.0, pchembl_grouping='combined')
```

**Option: Local mode**

```
from cpiextract import Comp2Prot
from cpiextract.utils import load_dbs, load_pubchem_files

data_path = 'data/Databases/'
dbs = load_dbs(data_path)
pubchem_files = load_pubchem_files(data_path)

C2P = Comp2Prot(execution_mode='local', dbs=dbs, pubchem_files=pubchem_files)

interactions, states, raw_interactions = C2P.comp_interactions(
    input_id='ZPEIMTDSQAKGNT-UHFFFAOYSA-N',
    pChEMBL_thres=3.0, strong_positive_thres=6.0, pchembl_grouping='combined')
```

Full documentation, local execution instructions, and further exemplary notebooks are provided in the https://github.com/menicgiulia/CPIExtract. The source code is publicly available under an MIT license on the GitHub.

### User scenarios

CPIExtract requires the user to download and preprocess the nine databases prior to running any pipeline. PubChem, ChEMBL, and OTP are exceptions as these databases have APIs. If these databases are not provided a local path and the user requests extraction from these databases, then CPIExtract will access them via their APIs. It should be noted that any reliance on APIs will greatly impact performance due to query limits within each API. We strongly recommend preprocessing the databases to obtain significantly faster execution times. To ease this process, we provide two notebooks to download (db_download.ipynb) and preprocess (db_preprocessing.ipynb) the databases on our GitHub https://github.com/menicgiulia/CPIExtract. Given the notable file size of these databases, we developed the package to support different modes of execution based on storage and software availability. Currently, there are two execution modalities: local and server.

#### Local execution

For this execution mode, the user needs to have downloaded the datasets or the portions of the datasets they are interested in. Note that if run with `interactions_select` functions, then the user will only need local copies of the select databases they want to query. The datasets are stored in CSV or TSV format and are loaded into the program environment as DataFrames using `pandas` Python library. The user can then initialize the pipeline classes, passing a dictionary containing the loaded DataFrames as parameter to the class constructor. The described mode is the fastest, as the files are loaded locally and readily available. However, the improved performance comes at the cost of increased storage and memory requirements. The combined databases occupy up to 13GB of disk space and are all loaded into the program simultaneously, thus demanding at least 16GB of RAM to run the package.

#### Server Execution

For server execution, the user does not need any database file stored locally. Instead, the pipeline connects to a remote MySQL server to retrieve raw data and store it in a DataFrame for the following filtering phase. When selecting this modality, the user passes the configuration to connect the MySQL server to the pipelines' constructors: address, user, password, and schema. The server is then accessed using the `mysql-connector` Python package, extracting raw data through select queries to the databases. This execution mode requires the setup of the MySQL server before executing the pipelines. Furthermore, the connection could represent a major bottleneck for the pipeline execution, as all raw data are transferred over the network before being processed in the remaining steps. On the other hand, it offers the advantage of being more scalable. It allows the use of the package on multiple machines with remote access to the same MySQL server, thus demanding less local storage.

#### APIs limitations and runtime performance

The PubChem API causes considerable performance limitations in the pipeline. Specifically, in the Prot2Comp pipeline, this API allows the search of a single compound per request when using external databases' identifiers as input. To speed up the data harmonization and extraction of interacting compounds data, PubChem's Compound Identifiers (CIDs) are desirable. This information is absent from most databases used by the package. To overcome this limitation, we preprocessed the downloaded databases by extracting CIDs and InChIKeys from the PubChem website and then merging this information using the database's external identifiers. We have provided the code to perform the preprocessing (db_preprocessing.ipynb) on our GitHub https://github.com/menicgiulia/CPIExtract for each database. We recommend performing this step before running the Prot2Comp pipeline, as it will allow users to collect data significantly faster. The amount of RAM required to complete this step is dependent on the initial dataset's size, for example, PubChem will require around 20GB RAM to complete while OTP will require less than 2GB RAM. If PubChem cannot be preprocessed, CPIExtract will fall back on API calls. Preprocessed datasets at publication are available on the GitHub (see SI: Database Information at Publication).

To demonstrate the runtime performance of CPIExtract under realistic usage conditions, we benchmark the package with both local and API modes using two different datasets based on the number of interactions collected. We randomly sampled 100 compounds with confirmed interaction data ($\geq$ 50 interactions across all databases) and 100 compounds confirmed to return no interactions, yielding 200 compounds in total. Each compound was queried using `Comp2Prot.comp_interactions()`. Note that in local mode all nine databases were queried from local files, and API calls are only made to PubChem and Biomart to unify the compound and protein identifiers. In API mode there are three databases which require API calls to query for interactions: PubChem, ChEMBL, and Open Targets Platform, in addition to the API calls to unify compound and protein identifiers. All other datasets do not have an API for queries, instead their datasets require a local copy to process within CPIExtract. These datasets remained as local in both modes for a balanced comparison between the modes. The same 200 InChIKeys were used across both modes to ensure direct comparability. Per-query time was recorded for each compound, and mean and standard deviation are reported stratified by interaction count and mode (Table S1). The harmonization of identifiers is performed by PubChem API (compounds) and biomart API (proteins). Both Local and API modes can be improved by using mygene instead of biomart for protein identifier harmonization. API mode runtimes reflect real world usage conditions including enforced rate-limiting delays inherent to each external service. All benchmarks were run on a single CPU core to reflect the default single-threaded execution of the package to meet query per unit time requirement of each API. Thus, the runtimes shown for local mode can be improved through parallelization whereas API mode cannot utilize parallelization. At publication, running all the preprocessed datasets in local mode requires at least 30GB RAM.

| **Mode Type** | **Interaction Count** | **Mean Time (s)** | **SD Time (s)** |
|---|---|---|---|
| Local | 0 | 6.1 | 0.70 |
| Local | $\geq 50$ | 15.7 | 17.0 |
| API | 0 | 22.3 | 22.2 |
| API | $\geq 50$ | 76.1 | 102.8 |

Table S1: Per-query runtime (mean ± SD, seconds) for CPIExtract in local and API execution modes, stratified by whether interaction data were found (0 vs. $\geq 50$ interactions). Results are based on 100 randomly sampled InChIKeys per stratum. API mode uses remote calls to PubChem, ChEMBL, and Open Targets Platform; all other databases remain local in both conditions. biomart API harmonizes protein identifiers.

### Database Information at Publication

We have provided preprocessed datasets for all the databases used with CPIExtract on the GitHub at https://github.com/menicgiulia/CPIExtract/tree/main/data. These datasets were downloaded in December 2025. Many of these resources require users to manually download the information from their websites though few have automated systems. At publication the GitHub contains the below databases and CPIExtract performs the following filtering operations:

- BindingDB - Downloadable via https://www.bindingdb.org/
  - Target Source Organism = Homo sapiens
  - Single protein targets (excludes complexes)
  - Excludes targets with point mutation(s)
  - Accepts temperature $<$ 40°C or missing (assume normality if missing)
  - Accepts $5 < \text{pH} < 9$ or missing (assume normality if missing)
  - pChEMBL value computed independently from each non-null activity column (Ki, IC50, Kd, EC50, all in nM)
  - Duplicate records removed
- CTD - Downloadable via https://ctdbase.org/
  - OrganismID = 9606 (Homo sapiens)
  - InteractionActions contains `affects^binding`, `increases^binding`, or `decreases^binding` (restricts to direct ligand–protein binding interactions)
  - GeneForms = protein (excludes non-protein gene products)
  - No activity value reported

- ChEMBL - Accessible via https://ftp.ebi.ac.uk/pub/databases/chembl/
  - Target Organism = Homo sapiens
  - Activity Comment not in {Not Determined, Not Evaluated, inconclusive, undetermined, No data}
  - Source ID = 1 (interactions extracted from scientific literature only)
  - Data Validity Comment is null or "Manually validated"
  - pChEMBL Value is not null
  - Target Type = SINGLE PROTEIN (excludes cell lines, protein complexes, RNA, etc.)
  - Duplicate records removed
- DTC - Downloadable via http://drugtargetcommons.fimm.fi/
  - `standard_type` restricted to a defined set of potency/activity measures (IC50, Ki, EC50, Kd, ED50, XC50, AC50, CC50, AVERAGEIC50, ACTIVITYEC50, SC50, AD50, DC50, XI50, GC50, GI50, and their log-transformed variants: LOGEC50, -LOGIC50, PIC50, LOGIC50, LOGKD, LOGKI, PEC50, PIC50(CALC), LOG(10^6/IC50), -LOGED50, LOG1/IC50, PED50, LOG1/KD, LOG(1/IC50), 1/KI, PXC50, PKI, PKD, PA2, PKB)
  - `standard_units` restricted to a defined set of convertible units (nM, M, /nM, nmol/L, mg.kg$^{-1}$, $\mu$g.mL$^{-1}$, $\mu$mol.kg$^{-1}$, ppm, mg/mL, ng mL$^{-1}$, mM, nmol/mg, and several scientific-notation variants), or missing when already in dimensionless pChEMBL scale
  - `standard_value`, `target_id`, and `compound_id` all non-null
  - `doc_type` non-null (interaction must have a reported source)
  - `activity_comment` is not inconclusive

  - mutated targets excluded: `wildtype_or_mutant` $\neq$ mutated and `mutation_info` is null
  - All values standardized to nM; zero and infinite values removed
  - Duplicate records removed
- DrugBank - Downloadable via https://go.drugbank.com/ (requires license)
  - Organism = Humans
  - Non-null DrugBank ID and HGNC identifier required
  - No quantitative activity value reported
- DrugCentral - Downloadable via https://drugcentral.org/
  - ORGANISM = Homo sapiens
  - ACT_TYPE $\in$ {IC50, Ki, EC50, Kd, AC50}
  - ACT_SOURCE $\neq$ UNKNOWN
  - ACTION_TYPE is non-null
  - ACTION_TYPE not in {PHARMACOLOGICAL CHAPERONE, RELEASING AGENT, ANTIBODY BINDING}
  - TARGET_CLASS not in {CD molecules, RNA, Unclassified, Viral envelope protein, Polyprotein}
  - ACT_VALUE is non-null and non-zero
  - pChEMBL value computed from ACT_VALUE
  - Duplicate records removed
- OTP - Accessible via http://ftp.ebi.ac.uk/pub/databases/opentargets/
  - Only mechanism-of-action (MoA) records retained; requires a non-null target name and at least one associated target identifier

  - targetType = single protein
  - Implicitly human-restricted (queried by human Ensembl gene ID / ChEMBL drug ID)
  - No quantitative activity value reported

- PubChem - Accessible via https://ftp.ncbi.nlm.nih.gov
  - Human targets only (Gene ID cross-referenced against `TaxonomyID = 9606`)
  - Activity Outcome not in {Unspecified, Inconclusive}
  - Activity Name $\neq$ CD
  - Activity Unit = $\mu$M or nM
  - Non-null, non-empty compound identifier and activity value

- Stitch - Downloadable via https://stitch-db.org/
  - Restricted to human protein–chemical interactions (source data pre-filtered to human on download)
  - Experimental evidence score $> 0$ (restricts to interactions with direct experimental support, excluding database, text-mining, or prediction evidence)
  - No quantitative activity value reported

All databases retain relational values and are used downstream to determine qualitative (presence/absence) records. If interaction has relational below user given pChEMBL threshold then CPIExtract outputs negative interaction. If above threshold then reports positive interaction. All negative statements such as "Inactive" and "Non-binding" are output as negative interactions.

In addition, we have provided code to automatically download and preprocess the current dataset versions where possible (db_preprocessing.ipynb) on our GitHub https://github.com/menicgiulia/C[illegible] for each database to prepare them for CPIExtract queries.

### Database selection

In addition to running the standard pipelines, users can interactively select from which sources to extract interaction data. This can be done if users want to exclude one or more data sources from the output. By doing so, only the selected databases will be queried, speeding up the pipeline execution at the cost of reducing the data extracted for each input compound or protein. It is important to note that applying a filter on the pChEMBL threshold on the pipelines' outputs will implicitly exclude any interaction data reported in CTD, DB, OTP, and STITCH databases. Furthermore, selecting only a single source is equivalent to extracting data directly from that database, without performing any integration. CPIExtract outputs both an aggregated dataframe containing average pchembl values for each interaction over all nine databases as well as a filtered dataframe that outputs each database's results without aggregation for traceability of the values used in the averaging.

### Optional Filtering Parameters

In addition to the base filtering steps described in the main paper, users can interactively perform additional filtering with optional parameters:

- `pChEMBL_thres`: Users can choose the threshold used to classify the strength of an interaction. This does not remove any interaction from the output - every interaction that passes each database's own filtering is retained regardless of its value. Instead, `pChEMBL_thres` determines whether a pair with quantitative evidence is labeled negative or positive for interaction, and gates which censored (inequality) measurements count as confirmed positive or negative evidence. (Default is 3.0).

- `strong_positive_thres`: For a pair classified as positive, this threshold splits the label further into `strong positive` (average pChEMBL above or equal to this value) or `weak positive` (above `pChEMBL_thres` but at or below this value). (Default is 6.0).

- `pchembl_grouping`: Controls how the average pChEMBL for a compound-protein pair is computed across its individual measurements. `'combined'` reports a single average across all measurement types together, alongside separate averages for binding/affinity-constant measurements (e.g. Ki, Kd) and 50%-response potency measurements (e.g. IC50, EC50), all in the same output row. `'unique'` instead reports a separate average per exact measurement type present in the data. (Default is `'combined'`).

- `experimental_thres`: The minimum STITCH/STRING experimental confidence score (0–999 scale) required for a Stitch interaction to be retained. (Default is 400, STRING's own documented low/medium-confidence boundary).

- `protein_types` (compound-input direction only): Restricts which of DrugBank's protein role categories - target, enzyme, carrier, transporter - are included. Defaults to all four if not specified.

For stereoisomer handling, every database is always searched using the input compound's full structural class, retrieving interactions across all of its stereoisomers rather than only the specific one submitted. Each returned interaction is labeled in a `structure_match` column as `stereochemical` (an exact match to the input stereoisomer) or `scaffold` (a match to the same base structure but a different stereoisomer), so users can filter to exact matches themselves if desired.

### Using CPIExtract in Computational Approaches

Naive train/test splits of compound–protein interaction data often yield artificially high predictive performance but poor generalizability[2], as AI/ML models learn shortcuts based on dataset annotation biases rather than underlying biological and chemical principles. This occurs when splitting compound–protein interaction data at kinetic-constant thresholds (e.g., on Ki, Kd, IC50, or EC50) to train ML/AI models on positive and negative pairs, or simply using presence/absence data, creating highly imbalanced datasets because of the sparse reporting of robust negative (non-binding) results within the scientific literature. Following the network-based sampling strategy introduced in AI-Bind[2], CPIExtract's harmonized, cross-source compound and protein identifiers can curate balanced training data by creating a bipartite compound–protein network, with compounds and proteins as nodes and strong positive interactions (e.g., pChEMBL $\geq 6$) as edges. Within this bipartite network, compounds and proteins separated by at least 7 hops are, on average, associated with weaker or absent binding evidence, providing a principled criterion for sampling additional, network-derived negative examples. Supplementing strong positive annotations with these network-derived negatives produces balanced training and test sets that prevent models from shortcut learning. CPIExtract's harmonized identifiers make this network construction, strong-positive classification, and hop-distance-based negative sampling directly reproducible from its own output, without the cross-source identifier reconciliation this process otherwise requires.

CPIExtract's output retains essential metadata that supports direct quality-based filtering of training data to ensure that computational models learn on high-quality information. For each compound–protein interaction, CPIExtract reports: (1) the number of source databases reporting the interaction, allowing users to prioritize pairs with broader cross-database consensus; (2) the binary presence/absence of the interaction in each individual source database, allowing users to include or exclude specific databases based on their own trust in a given source's curation standards; (3) the average pChEMBL value across contributing measurements, allowing filters by interaction strength; (4) the standard deviation of pChEMBL values across contributing measurements, allowing users to filter interactions by cross-source strength agreement; and (5) the specific measurement type(s) (Ki, Kd, IC50, EC50) contributing to each pair, allowing users to restrict training data to a single, pharmacologically consistent measurement type where mixing types is undesirable for a given modeling task. Together, these fields allow a user to construct a training set filtered to confidence thresholds appropriate to the application. This capability is complementary to, and can be combined with, network-derived negatives and other leakage-aware splitting approaches, for example PLINDER[1] and Bemis–Murcko scaffolds[3], to better generalize models.

### CPIExtract on Food Compounds

To explore the consistency of the compound-protein interactions within the queried food compounds, we assessed the coverage—how many of the sources report each interaction, and the consistency—how well activity affinities agree between sources. We find that the vast majority of compound–protein pairs (91.8%) are reported by only a single database within the nine queried by CPIExtract (Fig. S1a). The lack of overlap between the databases likely stems from the limited research on naturally occurring compound interactions, underscoring the value of CPIExtract in aggregating these data. By looking at the co-occurrence of each pair across the nine source databases, we see that the databases that report experimental binding affinities (PubChem, ChEMBL, BindingDB, and DTC) have the largest overlap, while the presence/absence databases show the most diversity in interaction data (Fig. S1b). 82.3% of pairs are found to have no pChEMBL value, while a further 11.3% are supported by exactly one pChEMBL value, showing that the majority of pairs rarely have a single pChEMBL value (Fig. S1c). These results combined indicate that the majority of compound-protein pairs with any overlap likely consist of pairs that contain heavily researched compounds and/or proteins within the literature. Therefore, when assessing consistency across databases, we can only use the 6.5% of the more heavily studied pairs with multiple independently reported pChEMBL values available. We report the conflict rate, defined as the proportion of pairs whose reported pChEMBL values differ by more than a given standard deviation threshold (Fig. S1d). For this assessment, we remove pairs with a standard deviation of exactly zero across contributing sources since these are likely instances of duplicate reporting (e.g., the same original measurement independently deposited into two databases) rather than true independent agreement. The conflict rate is found to increase with the number of independent quantitative sources at every threshold tested, from a mean of 24.6% among 2-source pairs to approximately 45.2% among 4-source pairs (>0.25 standard deviation threshold or 1.8x fold change). At >1.0 standard deviation (10x fold change), or one order of magnitude difference, this decreases to 5.6% among 2-source pairs and 12.7% among 4-source-pairs. Over the entire dataset (≥2-source pairs), we find the conflict rate is 7.8% at 1.0 standard deviation. These results show that, where binding affinities are present within the databases, they largely agree within the same order of magnitude, despite the aggregation of different binding affinity measurements (Ki, Kd, IC50, EC50). Indeed, the increase in conflict rate between 2-source pairs and 4-source pairs likely stems from the greater chance to have a higher standard deviation over the differing binding affinities, while lesser-cited pairs are reported with the same binding affinity measurement.

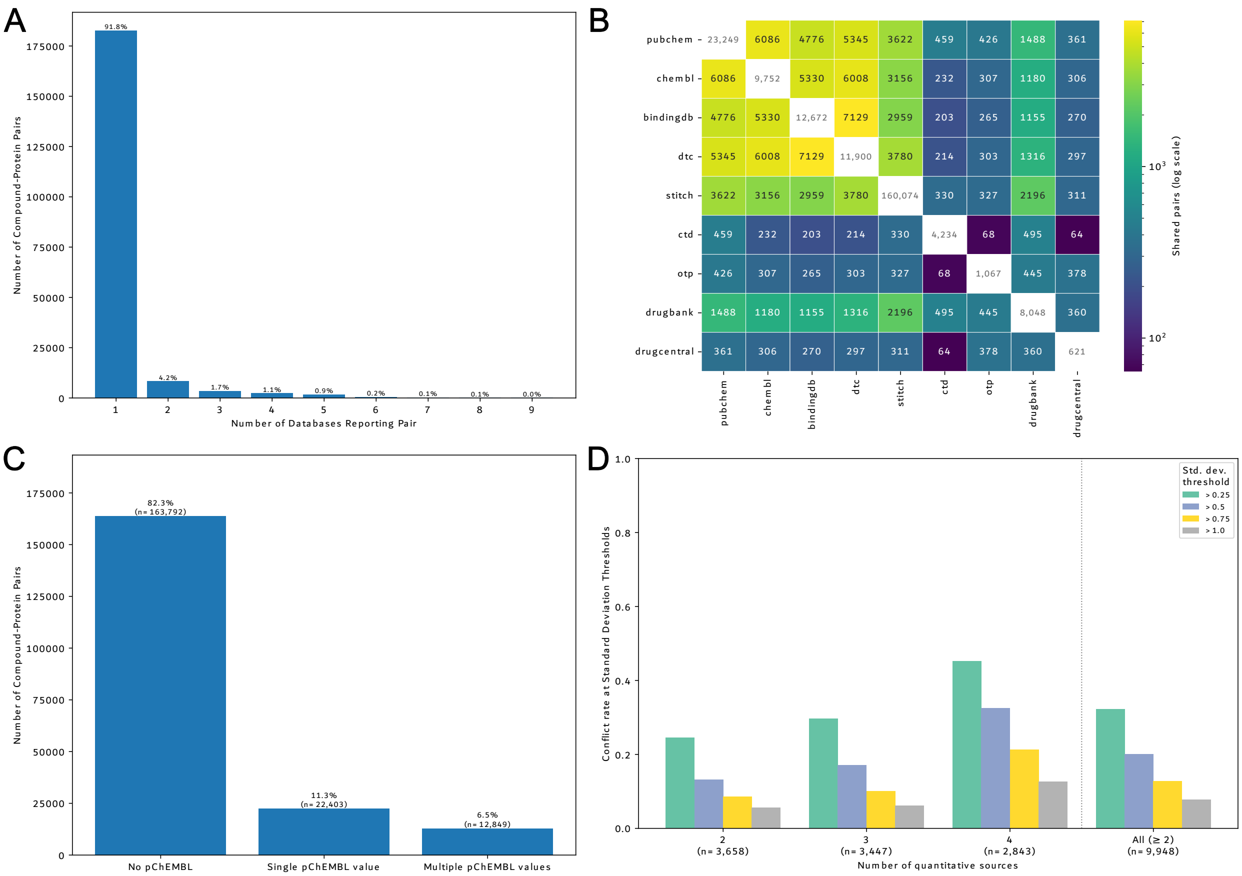


Figure S1: **Cross-source coverage and consistency assessment.** **A)** Distribution of compound-protein pairs by the number of reporting databases across all nine CPIExtract sources. **B)** Pairwise co-occurrence between the nine source databases, alongside each source's total individual coverage (diagonal). **C)** Composition of compound-protein pairs by number of contributing pChEMBL activity values, considering the four quantitative sources (PubChem, ChEMBL, BindingDB, DTC). **D)** Conflict rate calculated by the proportion of pairs whose pChEMBL values differ by more than a given standard deviation threshold among the pairs with exactly 2-sources, 3-sources, 4-sources, and All (≥2-sources).

### KEGG pathway Enrichment

To evaluate whether the additional compound–protein interactions recovered by CPIExtract translated into biologically meaningful pathway-level signal, we performed a KEGG pathway enrichment analysis comparing CPIExtract-derived interactions with curated DrugBank interactions.

For each compound, we independently built two target sets: one using CPIExtract and one using DrugBank. CPIExtract targets were obtained from the HGNC gene symbols associated with each compound in the CPIExtract output table. DrugBank targets were extracted from the DrugBank XML file, retaining human target proteins and their corresponding gene symbols. Compounds were matched between resources using full InChIKey identifiers, with the first InChIKey block used as a fallback structural match when necessary. For DrugBank compounds, DrugBank identifiers were also used to recover additional target annotations when available.

KEGG enrichment was performed independently for each compound and source using Enrichr through `gseapy`, with the `KEGG_2021_Human` gene set library. Enrichment was only computed for compound–source target sets containing at least five genes. For each enriched pathway, the adjusted $p$-value provided by Enrichr was used as the pathway significance measure. Pathways with adjusted $p < 0.05$ were considered significant.

For each compound–pathway pair, we quantified the enrichment signal as:

$$S = -\log_{10}(p_{\mathrm{adj}})$$

where $p_{\mathrm{adj}}$ denotes the adjusted enrichment $p$-value. Non-significant compound–pathway associations were assigned an enrichment signal of zero. The differential enrichment between CPIExtract and DrugBank was then computed as:

$$\Delta = -\log_{10}(p_{\mathrm{adj,CPIExtract}}) - (-\log_{10}(p_{\mathrm{adj,DrugBank}}))$$

Positive $\Delta$ values indicate stronger pathway enrichment when using CPIExtract-derived targets, whereas negative values indicate stronger enrichment when using DrugBank-derived targets.

#### ATC-based compound grouping

To further investigate whether the enrichment gains differed across compound classes, we defined a study compound panel composed of two groups. First, we selected compounds from

DrugBank annotated with Anatomical Therapeutic Chemical (ATC) codes corresponding to vitamins and general nutrients. Specifically, compounds with ATC prefix `A11` were classified as *vitamins*, and compounds with ATC prefix `V06` were classified as *general nutrients*. Together, these compounds represent the natural or endogenous-like compound group used to assess whether CPIExtract improves pathway coverage for compounds that are typically less represented in curated drug–target resources. Second, we included a set of therapeutic compounds previously evaluated in the present analysis. These compounds were labelled as *drug compounds* in the supplementary heatmap.

Thus, the final compound panel used for the supplementary differential KEGG enrichment visualization consisted of: (i) ATC `A11` vitamins, (ii) ATC `V06` general nutrients, and (iii) previously studied therapeutic drug compounds.

#### KEGG pathway category annotation

To facilitate biological interpretation of the enriched pathways, KEGG pathways were annotated using the official KEGG BRITE pathway hierarchy. The top-level KEGG categories considered were: Metabolism, Genetic Information Processing, Environmental Information Processing, Cellular Processes, Organismal Systems, Human Diseases, and Drug Development. Pathways that could not be mapped to the KEGG hierarchy were labelled as unmapped.

#### Differential enrichment heatmap

Supplementary Fig. S2 shows the differential KEGG enrichment profile between CPIExtract and DrugBank for the selected compound panel. Rows correspond to KEGG pathways and columns correspond to representative compounds. Cell values represent the differential enrichment score $\Delta$, defined above. Red values indicate stronger enrichment using CPIExtract-derived interactions, whereas blue values indicate stronger enrichment using DrugBank-derived interactions. For visualization purposes only, extreme values were capped at the 99th percentile of the absolute differential enrichment distribution, while the underlying differential enrichment matrix was left unchanged.

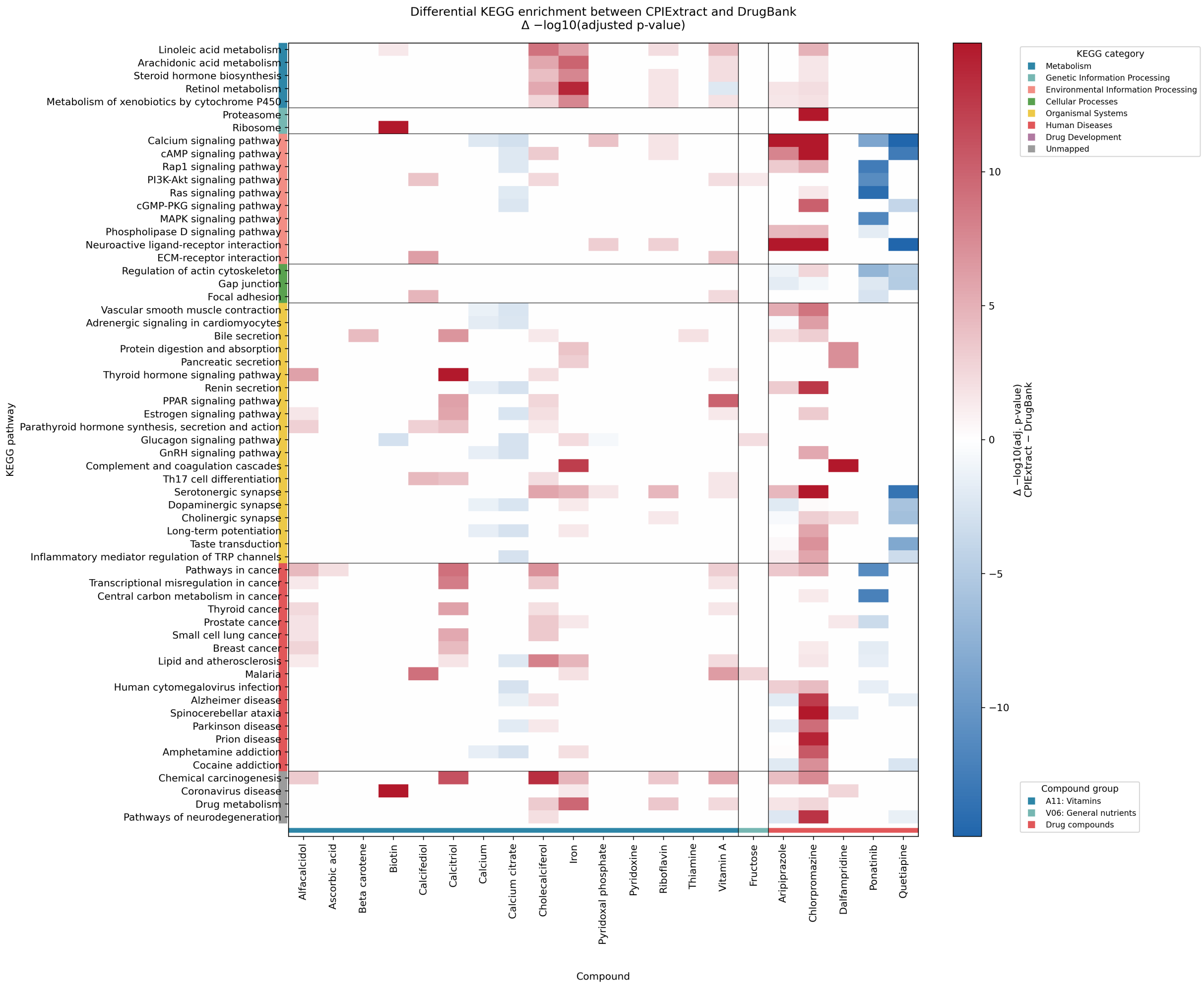


Figure S2: **Differential KEGG pathway enrichment between CPIExtract and DrugBank.** Each cell represents the difference in pathway enrichment significance between CPIExtract and DrugBank for a given compound–pathway pair, computed as $\Delta = -\log_{10}(p_{\text{adj,CPIExtract}}) - (-\log_{10}(p_{\text{adj,DrugBank}}))$. Positive values indicate stronger enrichment using CPIExtract-derived protein interactions, whereas negative values indicate stronger enrichment using DrugBank-derived interactions. Pathways are grouped according to the official KEGG categories. Compounds are grouped into ATC `A11` vitamins, ATC `V06` general nutrients, and previously studied therapeutic drug compounds.

This analysis illustrates that CPIExtract provides stronger pathway-level signal for many vitamin and nutrient compounds, especially across metabolic and signalling pathways. In contrast, therapeutic drug compounds generally show smaller differences between CPIExtract and DrugBank, and in some cases stronger enrichment using DrugBank-derived targets. These results support the observation reported in the main text that the additional CPIExtract interactions are particularly informative for naturally occurring or nutritional compounds whose bioactivity is less comprehensively represented in curated drug–target databases.

#### Conditional permutation controls

To assess whether the enrichment differences described above exceeded those expected under random target composition while retaining the observed target-set sizes, we performed additional conditional gene-label permutation analyses. These controls used a paired cohort and a common local enrichment procedure, as specified below.

**Paired cohort and source ablation.** We compared DrugBank, full CPIExtract, and CPIExtract after removal of associations supported exclusively by DrugBank. Associations with support from at least one non-DrugBank source were retained, including associations also reported by DrugBank. Gene symbols were deduplicated within each compound after filtering. Compounds were matched to DrugBank by exact full InChIKey, using the CPIExtract `db_inchikey` when available and `pc_inchikey` otherwise. DrugBank records sharing an InChIKey were merged by taking the union of their target sets. The paired cohort comprised 244 compounds with at least five distinct target genes in each of the three resources. DrugBank target sets were retained without intersection with CPIExtract targets. This supplementary analysis used all targets without a pChEMBL cutoff.

**Enrichment procedure and statistics.** We used a fixed snapshot of the `KEGG_2021_Human` gene-set library and a common background of 9,871 gene symbols, defined as the union of valid CPIExtract and DrugBank target symbols before compound matching. Pathway membership was restricted to this background; genes without KEGG annotation remained in the background and target sets. One-sided hypergeometric overrepresentation tests were performed for all 320 pathways, followed by Benjamini–Hochberg (BH) adjustment separately for each compound and resource. Zero-overlap pathways were included with $p = 1$. We evaluated two statistics: the number of significant pathways ($p_{\text{adj}} \leq 0.05$; primary statistic), and the pathway-averaged score using the transformation defined above, without setting nonsignificant scores to zero (secondary statistic),

$$A_{ir} = \frac{1}{320} \sum_{k=1}^{320} -\log_{10}\left(p_{irk}^{\text{adj}}\right), \qquad T_r = \frac{1}{244} \sum_{i=1}^{244} \left(A_{ir} - A_{i,\text{DrugBank}}\right). \tag{1}$$

Here $i$, $r$, and $k$ denote compound, resource, and pathway, respectively; $A_{ir}$ is the untruncated pathway-averaged score, distinct from the visualization score $S$ defined above. The analogous paired statistic was computed for significant-pathway counts. Each compound received equal weight. Figure S3 displays the secondary statistic for each resource relative to its own null model; Figure S4 displays the corresponding paired differences from DrugBank.

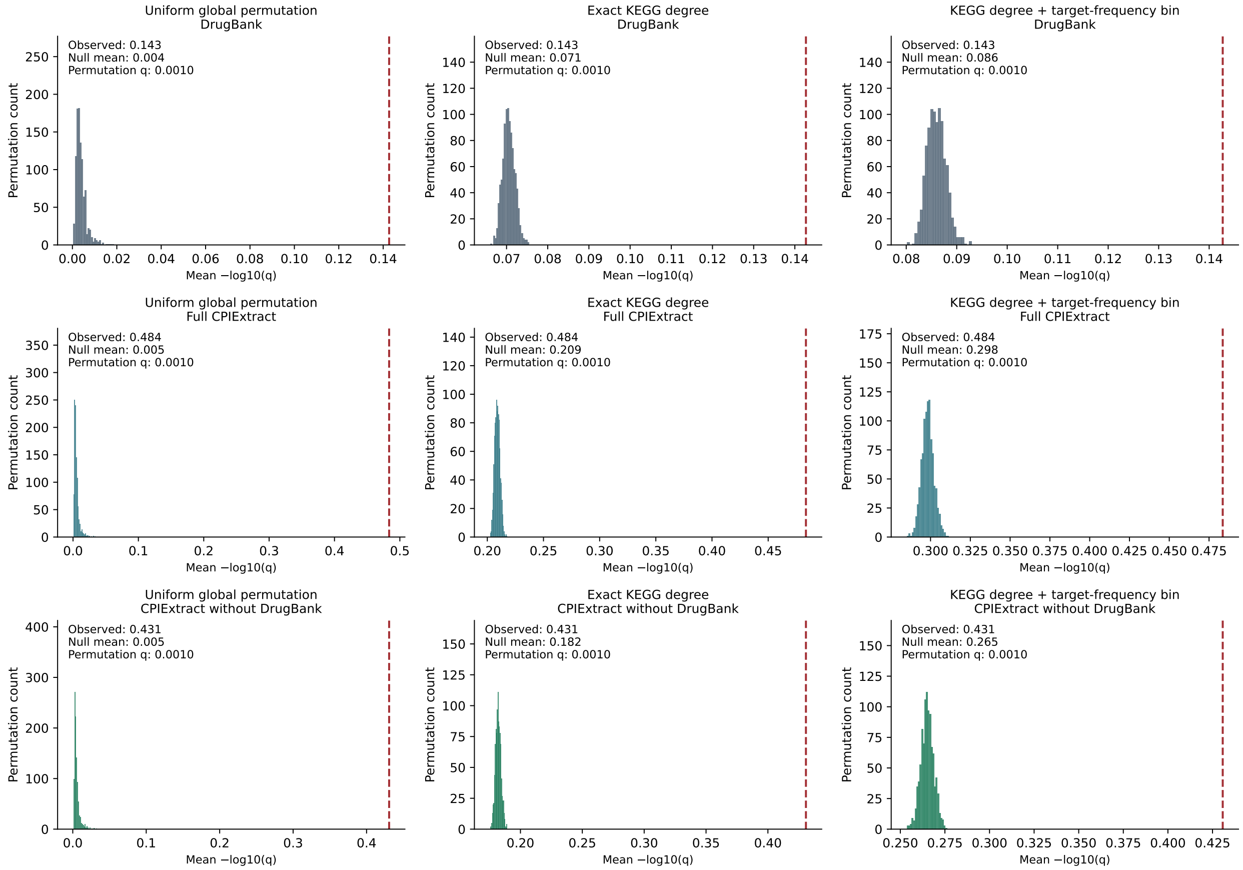


Figure S3: **KEGG enrichment signal for each resource relative to global conditional null models.** Histograms show 1,000 null values of the mean pathway-averaged $-\log_{10}(p_{\text{adj}})$ across the same 244 compounds. Rows show DrugBank, full CPIExtract, and CPIExtract retaining associations with non-DrugBank support. Columns show unrestricted global gene-label permutation, permutation within exact KEGG annotation-degree strata, and permutation within joint annotation-degree and target-frequency-bin strata. Each repetition applies one common gene-label reassignment to all compounds and resources, preserving target-set sizes and all overlaps. Red dashed lines mark observed statistics. Annotations report the observed statistic, null mean, and upper-tail permutation $q$-value, BH-adjusted across the 18 source-level tests (three resources, three null models, and two statistics). These permutation $q$-values are distinct from the adjusted enrichment $p$-values entering the score. Axis scales differ between panels. Departure of each resource from its own null model does not itself establish a difference between resources; that comparison is tested in Figure S4.

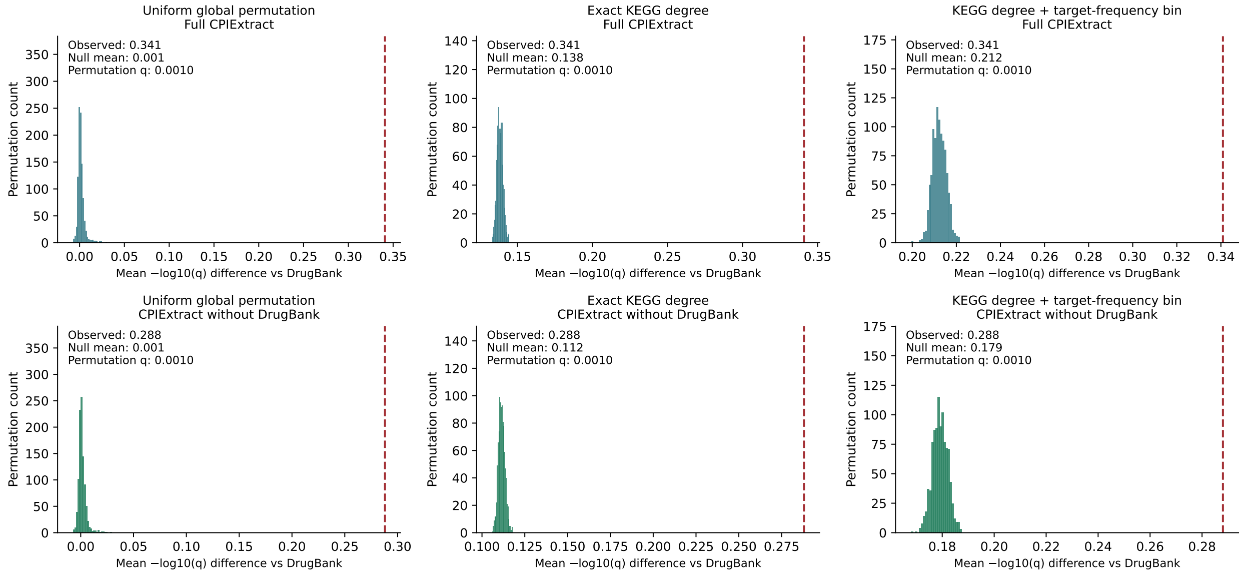


Figure S4: **Global conditional permutation controls for paired KEGG enrichment differences.** Histograms show 1,000 null values of the mean paired difference in pathway-averaged $-\log_{10}(p_{\mathrm{adj}})$ between CPIExtract and DrugBank across the same 244 compounds. The upper row uses full CPIExtract; the lower row retains associations with non-DrugBank support. Columns show unrestricted global gene-label permutation, permutation within exact KEGG annotation-degree strata, and permutation within joint annotation-degree and target-frequency-bin strata. Each repetition preserves all target-set sizes and overlaps across compounds and resources. Red dashed lines mark observed statistics. Annotations report the observed statistic, null mean, and BH-adjusted upper-tail permutation $q$-value; the latter is distinct from the adjusted enrichment $p$-values entering the score. Each histogram represents cohort-level statistics, rather than individual compound–pathway scores. Axis scales differ between panels.

**Global conditional null models.** For each of three null models, we generated 1,000 permutations using seed 20260907. Each repetition used one bijective reassignment of the 9,871 background gene labels, applied jointly to all compounds and resources. This preserved every target-set size and all within- and between-compound overlaps, including overlaps between resources. The models allowed exchanges (i) among all background genes, (ii) only among genes annotated to exactly the same number of KEGG pathways, or (iii) only among genes with both the same exact KEGG annotation degree and the same pooled target-frequency bin (0, 1, 2–3, 4–7, 8–15, and successive powers-of-two intervals). Frequency counted unique compound–gene pairs in the union of full CPIExtract and DrugBank before cohort selection. It measures representation in the supplied data, rather than experimental evidence quality. Genes in singleton strata remained fixed. The enrichment tests, BH adjustments, and aggregate paired statistics were recomputed for each permutation.

**Permutation inference and results.** Upper-tail empirical permutation $p$-values were calculated as $(b+1)/(1000+1)$, where $b$ is the number of permuted statistics at least as large as the observed statistic, with conservative numerical tie handling. BH correction was applied jointly to the 12 aggregate paired tests (three models, two CPIExtract variants, and two statistics). The 18 source-level tests were adjusted as a separate family. For the secondary statistic, the observed paired differences were 0.34091 for full CPIExtract and 0.28804 after removal of DrugBank-only evidence. Under the most conditioned null, the corresponding null means were 0.21229 and 0.17915, with central 95% null intervals [0.20657, 0.21830] and [0.17340, 0.18487]. The excesses over the null means were therefore 0.12862 and 0.10888. All six upper-tail paired tests for this statistic had empirical $p = 1/1001$ and BH-adjusted permutation $q = 0.000999$. For the primary statistic, the observed mean differences were 15.53 and 12.77 significant pathways, compared with null means of 12.62 and 10.54 under the most conditioned model, giving excesses of 2.91 and 2.24 pathways, respectively, with the same empirical significance.

These results demonstrate departure from the specified conditional random-composition models while retaining complete target sets. They do not establish superior enrichment at equal target counts or greater interaction precision. The background is a pooled observed-target universe rather than the complete human proteome. Conditional exchanges do not control all possible biases, and source ablation cannot exclude indirect reuse of DrugBank evidence by other databases.

### Comp2Prot on DB

To further study the performance of CPIExtract, we performed additional evaluations and comparisons on the data produced by the Comp2Prot pipeline. We studied the quality of all the interactions extracted from the data sources using the 3 annotation classes described in the main paper: non-binding (pChEMBL$\leq$ 3), weakly binding (3 $<$pChEMBL$<$ 6), and strongly binding (pChEMBL$\geq$ 6) (Fig. S5A). STITCH provided most of the interactions with missing bioactivity, while BDB, DTC and PubChem were the main sources of weakly and strongly binding annotations. Using this additional information, we could assign bioactivity data to the DB interactions for which DB does not provide any. Specifically, we extracted binding information for more than 29.1% of DB annotations, of which 17.8% were defined as strongly binding (Fig. S5B). We also evaluated the information gain compared to DB by selecting some DB candidates and measuring the $k_i$ ratio between CPIExtract and DB at different pChEMBL thresholds (Fig. S5C). For each candidate, the ratio showed generally more annotations collected. For some drugs, at higher thresholds, the ratio decreased to 1 ($k_i$(CPIExtract) = $k_i$(DB)). It is worth noting that the drop from no threshold to pChEMBL$\geq$ 0 is caused by the loss of interactions from STITCH and CTD which lack bioactivity data. To prove the necessity of our tool, we also examined the annotation overlap of databases with different pChEMBL thresholds (Fig. S5D). Most annotations were found in only one source, and less than 10% of annotations were reported in five databases or more. This implies again that most interactions are reported in only a few databases due to the significant heterogeneity in the types of information stored.

#### DrugBank Candidates Selection for Evaluation

For our evaluation of the information gain obtained from the package, we evaluated the $k_i$ ratio between CPIExtract and DB for a set of selected candidates extracted directly from DB (Fig. S5C). The selection process was carried out to obtain the best compounds for

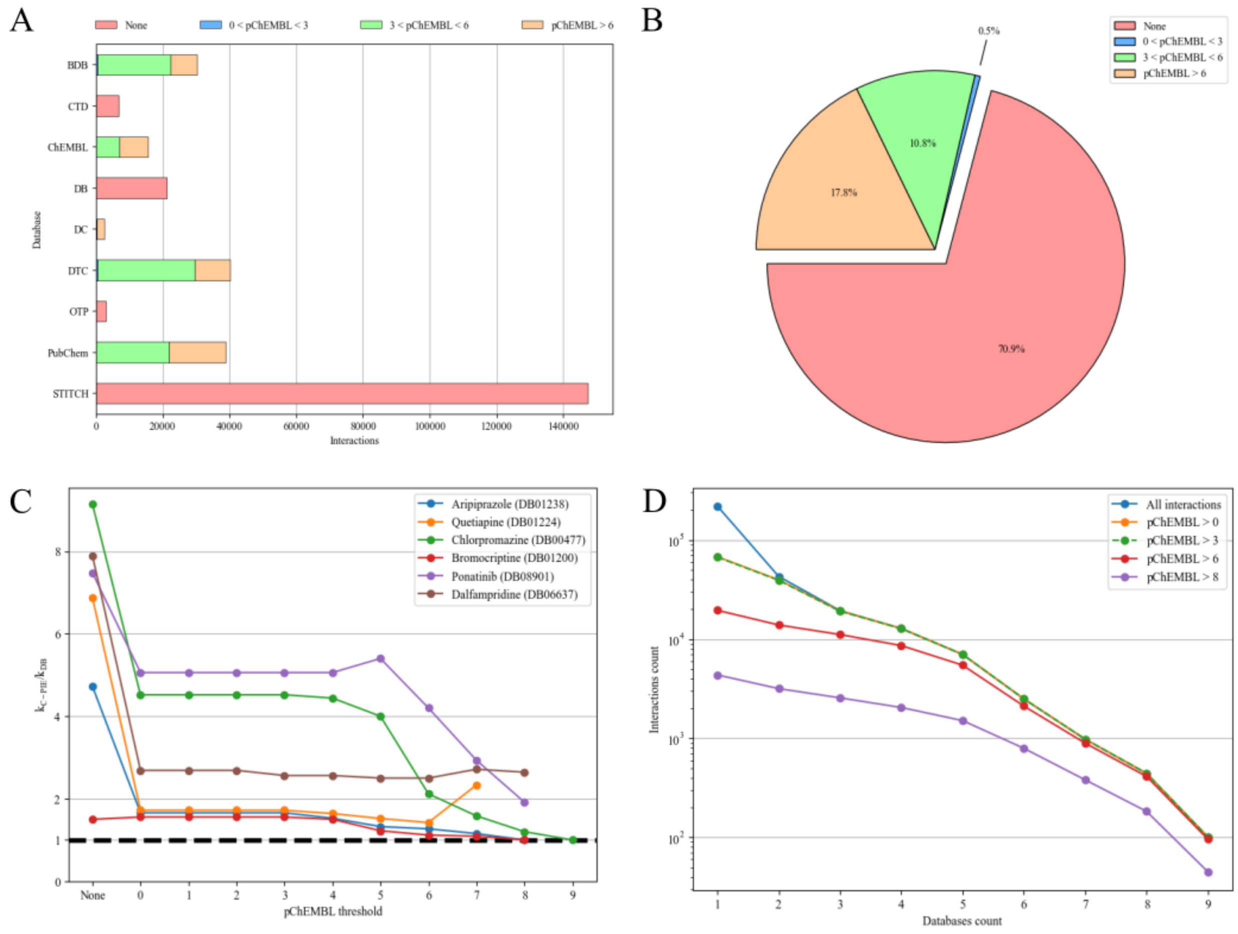


Figure S5: **Comp2Prot additional evaluation.** **A)** Distribution of no bioactivity data, non-binding, weak binding, and strongly binding annotations extracted from each database using the DB compound list as input. **B)** Percentages of no bioactivity data, non-binding, weak binding, and strongly binding DB interactions. Bioactivity data were extracted with CPIExtract from other databases. **C)** $k_i$ ratio between CPIExtract and DB at different pChEMBL thresholds for selected candidates. CPIExtract has more interactions per compound than DB. $k_i(\text{CPIExtract})/k_i(\text{DB}) = 1$ represents an equal number of interactions reported. **D)** Overlap of databases for interactions with different pChEMBL thresholds. The percentage of interactions reported in more than five databases is less than 10%.

which CPIExtract managed to obtain the most bioactivity data. Furthermore, we avoided compounds mostly interacting with CYP enzymes, carriers, and transporter (CCT) targets. This was done to exclude the metabolization drug interactions and focus more on those that caused health effects on the target. We curated the drugs following:

1. Given that DB does not provide bioactivity data, we matched DB interactions for each compound to those obtained by the package in order to extract pChEMBL values from

its output;

2. We computed the ratio between annotations with bioactivity data and the total annotations reported by DB;

3. We also computed the ratio of the annotations, including CCT targets, by the total annotations reported by DB;

4. We removed compounds with less than 15 unique targets, less than 80% of annotations with bioactivity data, and more than 50% of CCT targets.

This process produced the 6 compound candidates we used in our evaluation: Aripiprazole, Bromocriptine, Chlorpromazine, Dalfampridine, Ponatinib, and Quetiapine (Table S2).

| **Compound** | **ID** | **Targets** | **Bioactivity Ratio** | **CCT Ratio** |
|---|---|---|---|---|
| Aripiprazole | DB01238 | 38 | 0.8837 | 0.1163 |
| Quetiapine | DB01224 | 25 | 0.8333 | 0.2 |
| Chlorpromazine | DB00477 | 25 | 0.8621 | 0.2414 |
| Bromocriptine | DB01200 | 18 | 0.9 | 0.1 |
| Ponatinib | DB08901 | 17 | 0.8095 | 0.2857 |
| Dalfampridine | DB06637 | 16 | 0.8889 | 0.1111 |

Table S2: DB Candidates resulting from the selection process with the respective computed data. The candidates are sorted by number of targets.

### Prot2Comp on DB

We evaluated the Prot2Comp pipeline on the DB protein list. The pipeline collected and harmonized 3,293,463 unique interactions for 2,972 proteins. DB accounted for less than 1% of the total interactions found (Fig. S6A). Most of the data were extracted from BDB, ChEMBL, DTC, PubChem, and STITCH. Moreover, CPIExtract provided annotations for an additional 211 proteins which had no interaction data in DB. Next, we compared the quality of DrugBank interactions with CPIExtract, using pChEMBL as the evaluation metric (Fig. S6B). Prot2Comp provided 10 to 100 times more interactions for every pChEMBL threshold. Furthermore, it extracted many interactions not present in DB. For reference, the percentage of DB annotations for which Prot2Comp obtained bioactivity data was 28.9%, with the majority of them (17.6%) being with a pChEMBL$\geq$ 6 value (Fig. S6C).

There is a slight difference between the percentage of DB interaction strengths for the two pipelines that arises from occasional inconsistencies in DB. Specifically, for the same drug, DB reports an InChIKey that may differ from the one provided by the more authoritative source PubChem, used by the Prot2Comp pipeline to extract compound data for interactions. This, in turn, can cause the same drug-target pair to have a different number of sources. In Table S3, we provide an example between the drug Lestaurtinib (DB06469) and Receptor-type tyrosine-protein kinase FLT3 with both Comp2Prot and Prot2Comp. We see that the first block of the InChIKey (UIARLYUEJFELEN) is the same, indicating that the two entries are structurally identical; however, the second block of the InChIKey is different, revealing stereochemical differences as well as directionality differences between the two pipelines. PubChem harmonizes stereochemical issues between datasets and in fact links DB06469 to UIARLYUEJFELEN-LROUJFHJSA-N. Meanwhile, DrugBank reports its own unique stereochemistry. The drug-target pair is present in both pipelines. CPIExtract uses the stereochemistry as reported in the databases themselves, especially since many biological reactions are indeed stereochemistry-specific. In order to account for these differences, CPIExtract reports both InChIKey first block and full InChIKey as well as a "structure match" output column, so users can combine the various stereoisomers if desired.

| | **Comp2Prot** | **Prot2Comp** |
|---|---|---|
| **Compound DB ID** | DB06469 | DB06469 |
| **HGNC Symbol** | FLT3 | FLT3 |
| **InChIKey** | ***UIARLYUEJFELEN-LROUJFHJSA-N*** | ***UIARLYUEJFELEN-DMVVYWCZSA-N*** |
| **Avg. pChEMBL** | 8.535 | N/A |
| **pChEMBL Count** | 21 | 0 |
| **Sources Count** | 7 | 1 |

Table S3: Output difference for Comp2Prot and Prot2Comp for the drug-target pair Lestaurtinib and FLT3. The reported extracted InChIKey is different.

To further prove the information gain, we tested the degree $k_i$ (defined as the number of unique annotations for each protein $\text{prot}_i$) probability distribution of DB and CPIExtract using the thresholds pChEMBL$\geq$ 6 and pChEMBL $\geq$ 8 (Fig. S6D). In this case, the difference between DB and CPIExtract was more pronounced for both thresholds. Even when considering the higher threshold pChEMBL$\geq$ 8, the distribution had a significantly longer tail than DB. This is likely due to the narrow scope of DrugBank towards drug compounds, whereas by collecting data from various sources CPIExtract contains interactions for any compound be it a pharmaceutical, natural, or industrial compound. We evaluated the distribution of the $k_i$ ratio between CPIExtract and DB at different pChEMBL thresholds (Fig. S6E). On average, CPIExtract produced a huge information gain for every protein, even for higher pChEMBL thresholds. The information gain was even greater when excluding annotations with no bioactivity data reported. Finally, we show the importance of our tool by examining the annotation overlap of databases with different pChEMBL thresholds (Fig. S6F). Even for Prot2Comp, most annotations were found in only one source, with a small percentage being reported in five databases or more.

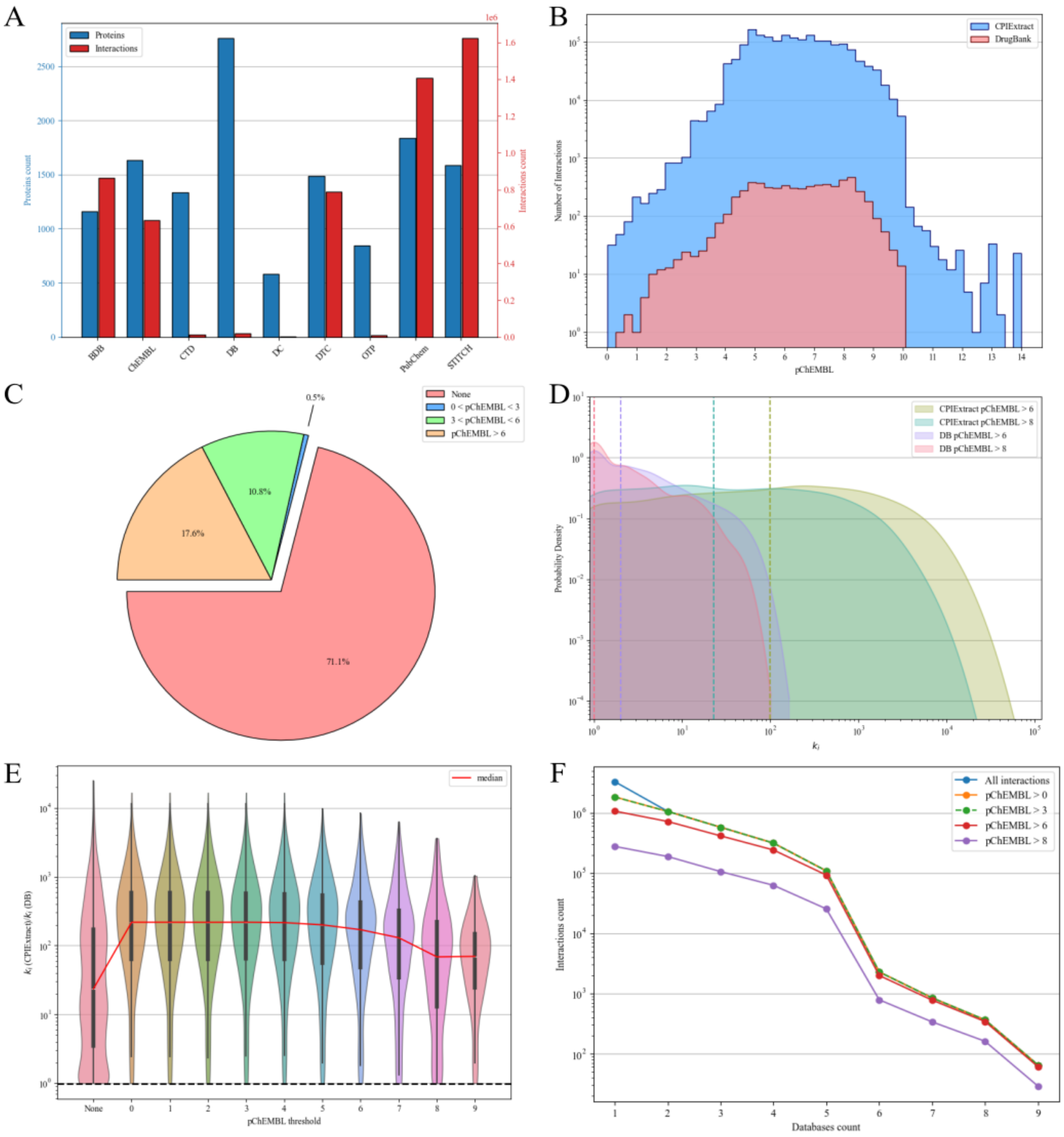


Figure S6: **Prot2Comp evaluation. A)** Databases' number of DB proteins for which interaction data was available and relative total annotation count. **B)** CPIExtract and DB interaction distribution over different pChEMBL values. DB data contains fewer interactions for each value and no interactions with pChEMBL$\geq$ 10. **C)**Percentages of no bioactivity data, non-binding, weakly binding, and strongly binding DB interactions. **D)** Density distribution of $k_i$ for CPIExtract and DB data at different pChEMBL thresholds. CPIExtract has more interactions per protein than DB, even with stronger thresholds. **E)** Distribution of $k_i$ ratio between CPIExtract and DB at different pChEMBL thresholds. $k_i(\text{CPIExtract})/k_i(\text{DB}) = 1$ represents an equal number of interactions reported. **F)** Overlap of databases for interactions with different pChEMBL thresholds. The percentage of interactions reported in more than five databases is around 10%.

### Comp2Prot on Naturally Occurring Compounds

CPIExtract can be used beyond pharmaceuticals to capture the biochemical interactions of compounds produced by industry, e.g., pesticides, or other compounds within the exposome. The food compounds within our diet are an essential component of the exposome. Here, we run the Comp2Prot pipeline on a list of naturally occurring compounds extracted from FooDB (`www.foodb.ca`), one of the largest resources on food chemicals. Specifically, we tested 56,552 compounds and compared the CPIExtract output with the same output for DB drugs. The pipeline collected and harmonized 73,260 interactions for 2,687 unique compounds (Fig. S7A). We evaluated the strength of the extracted data using pChEMBL values and compared it with the DB data (Fig. S7B). For FooDB compounds, CPIExtract extraction provided a similar number of non-binding and weakly binding interactions, whereas it provided on average ten times more strongly binding annotations for the DB drugs. The extracted bioactivity data provided quality information for 26.3% of all reported interactions, with 5.1% strongly binding, 20.2% weakly binding, and 1% non-binding annotations. We then compared the degree $k_i$ density distribution of both CPIExtract outputs for pChEMBL$\geq$ 6 and pChEMBL$\geq$8 thresholds (Fig. S7C). The degree distributions for each threshold proved to have the same median. Yet, DB drugs were shown to have, on average, more interactions than FooDB compounds, as proved by the higher probability density of the respective distributions for greater values of $k_i$. Finally, we evaluated the annotations overlap between CPIExtract databases (Fig. S7D) over increasing pChEMBL thresholds. The trend was similar to other CPIExtract results, proving that interactions are mainly found in a few data sources. Specifically, more than 90% annotations were found in three databases or less.

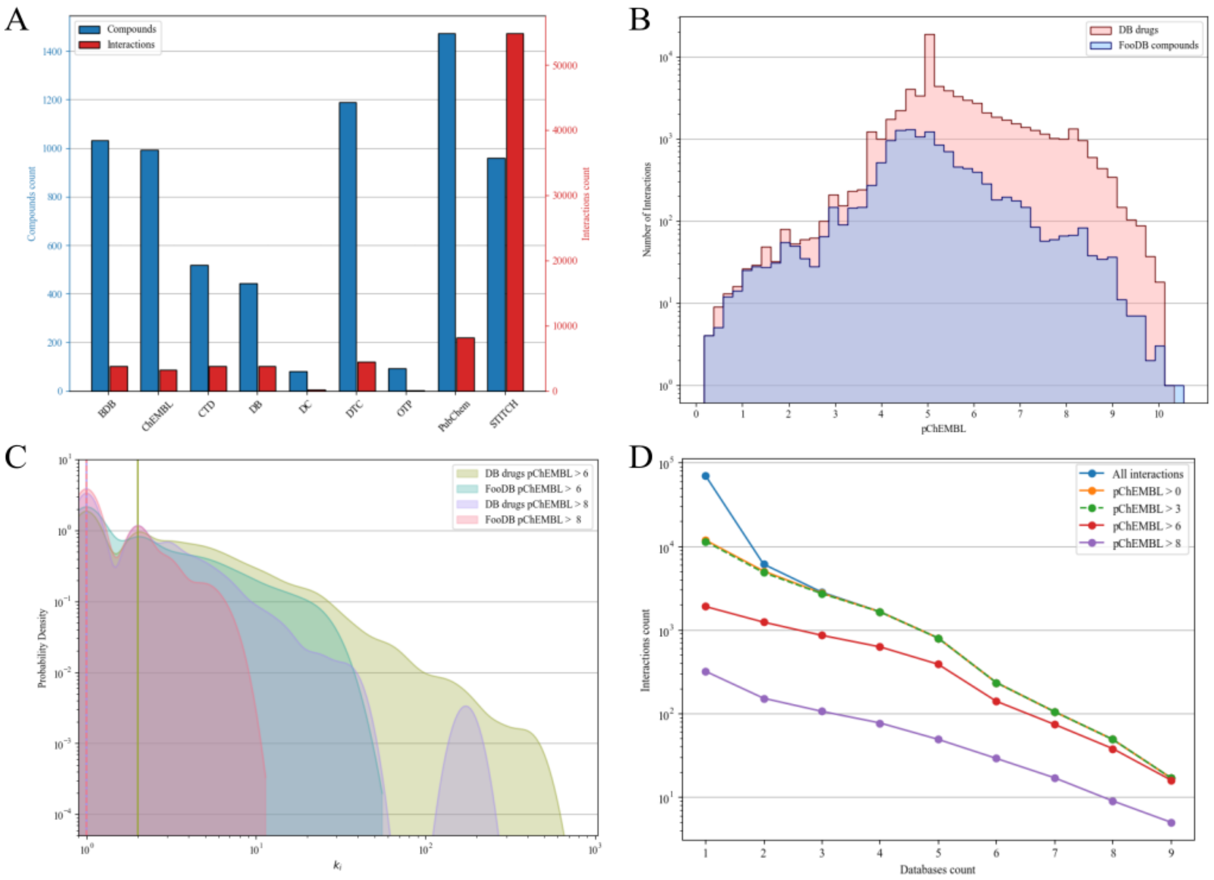


Figure S7: **FooDB compounds evaluation. A)** Databases' number of compounds for which interaction data was available and relative total annotation count. **B)** CPIExtract interaction distribution over different pChEMBL values for DB drugs and FooDB compounds. CPIExtract retrieved more interactions for DB molecules at higher pChEMBL thresholds. **C)** Distribution of $k_i$ for DB drugs and FooDB compounds data filtered using different pChEMBL thresholds. **D)** Overlap of databases for interactions with different pChEMBL thresholds. The percentage of interactions reported in less than three databases is 90%.
